# *Cryptococcus gattii* responds to mycobacterial exposure through coordinated remodelling of population dynamics and cell surface architecture, enhancing pulmonary persistence in a co-infection model

**DOI:** 10.64898/2026.05.25.727575

**Authors:** Hilaire Irere, Zar Ni San, Liliane Mukaremera, Ivy M. Dambuza

## Abstract

*Cryptococcus neoformans* and *Cryptococcus gattii* are major causes of fungal pneumonia and meningitis, frequently co-occurring with *Mycobacterium tuberculosis* in endemic regions, where co-infection is associated with increased mortality. Yet, how *Cryptococcus* adapts to mycobacterial co-presence within the lung remains poorly understood. Here, we show that mycobacterial cues trigger a conserved adaptive programme in *C. gattii*, mirroring responses previously observed in *C. neoformans*. Increasing exposure to mycobacteria drives cell and capsule enlargement and promotes titan cell formation, accompanied by dose-dependent remodelling of chitin and chitosan. Importantly, *in vivo* exposure to heat-killed mycobacteria increases *C. gattii* pulmonary burden, linking structural remodelling to enhanced persistence. These findings identify mycobacterial co-presence as a driver of fungal phenotypic plasticity and reveal pathogen-pathogen interactions as critical regulators of disease outcome, highlighting a previously unrecognised axis of co-infection relevant to *C. gattii* pathogenesis and therapeutic strategy.

## Introduction

Tuberculosis and cryptococcal disease represent two of the most significant infectious threats to global health. *Mycobacterium tuberculosis* (Mtb) causes an estimated 10 million new infections and approximately 1.3 million deaths annually, remaining one of the leading infectious causes of death worldwide^1^. Cryptococcal disease, driven primarily by *Cryptococcus neoformans* and *C. gattii*, causes over 150,000 deaths each year, predominantly due to cryptococcal meningitis in immunocompromised individuals. These microbes are designated World Health Organization priority pathogens owing to their high disease burden, clinical severity and increasing antimicrobial drug resistance.

*Cryptococcus* species are environmental basidiomycete fungi that cause life-threatening disease following inhalation of infectious propagules into the lung^2,3^. Among these, *Cryptococcus neoformans* and *Cryptococcus gattii* account for the vast majority of human cryptococcal infections worldwide^4,5^. After inhalation, the fungus establishes infection within the pulmonary environment^6,7^, a complex and heterogeneous niche shaped by host immune responses, nutrient limitation, inflammatory stress, and localized hypoxia that emerges during infection^8–11^. Successful persistence therefore requires fungal strategies that enable adaptation to these dynamic conditions. This adaptive flexibility is underpinned by several key virulence-associated traits that work in concert to ensure survival within the hostile host environment. For example, the polysaccharide capsule and the underlying fungal cell wall regulate exposure of immunostimulatory ligands, resistance to host-derived stresses, and survival within phagocytic cells^12–16^.

In addition, *Cryptococcus* display pronounced morphological heterogeneity, producing polyploid titan cells (≥10µm in diameter), typical yeast cells (∼5-7 µm), and smaller morphotypes (<5 µm)^17,18^. These cell types interact differently with host immunity and are increasingly recognized as an adaptive strategy that promotes persistence and dissemination during infection rather than representing stochastic variation^17–20^. While these virulence mechanisms have been extensively characterized in *C. neoformans*, *C. gattii* exhibits several distinctive features that set it apart from its more widely studied sibling. *C. gattii* presents a distinct and clinically important biology within this framework, fundamentally distinguished by its morphological profile and clinical presentation. Morphologically, *C. gattii* exhibits enhanced titan cell formation (>15 μm) with fewer small cell morphotypes compared to *C. neoformans*^21,22^. This bias toward enlarged cells has profound clinical implications: while *C. neoformans* more frequently affects immunocompromised hosts and readily disseminates to the CNS, *C. gattii* causes disease in immunocompetent individuals and shows a marked predilection for establishing persistent pulmonary infections, including cryptococcomas and prolonged lung residence^23–26^. The increase titanisation may mechanistically explain this clinical pattern, as larger cells are better equipped to resist lung environmental stresses but are less capable of crossing tissue barriers for dissemination. Although CNS infection can occur, extrapulmonary spread remains comparatively infrequent and poorly understood, suggesting that additional unknown factors may govern escape from the pulmonary niche^24^. Supporting this, *C. gattii* infections are associated with suboptimal protective immune responses in the lung, including impaired Th1 and Th17 polarization, suggesting that the fungus may be modulating host immune response, favouring long-term pulmonary colonisation^27^.

In its natural habits, *C. gattii* exists within complex microbial communities including soil and tree bark^28,29^, where it encounters diverse bacterial neighbours and environmental predators ^30–33^. Increasing evidence indicates that such cross-kingdom interactions can shape fungal physiology and virulence-associated traits^34–36^. Indeed, several studies have shown that bacterial molecules influence fungal morphology, capsule architecture, and cell wall composition in several fungal pathogens ^30,31,37^. Despite this growing recognition, the role of microbial interactions in shaping *C. gatti* biology remains poorly understood. The lung represents an ideal niche for such interactions, as both *M. tuberculosis* and *C. gattii* establish their primary infections within this shared pulmonary environment. Both pathogens are acquired via inhalation, establish residence within the alveolar space, interact extensively with macrophages, and can contribute to granulomatous inflammation^8,38–40^. Clinically, cryptococcal-tuberculosis co-infection is increasingly reported and is associated with poorer clinical outcomes and higher mortality rates^41,42^. However, despite overlapping epidemiology and host immune responses, the microbiology outcome of pathogen-pathogen in a co-infection framework remains relatively underexplored ^43,44^. We previously demonstrated that *C. neoformans* responds to mycobacterial exposure by reconfiguring morphological diversity and capsule architecture^45^. Whether *C. gattii* actively responds to the presence of *M. tuberculosis* and how such interactions might influence fungal adaptation or persistence, remains largely unknown. Using a physiologically relevant human plasma-like medium (HPLM), we examined the impact of heat-killed *M. tuberculosis* on fungal population structure, capsule thickness, and cell wall composition. Our findings reveal that mycobacterial-derived signals drive coordinated restructuring of *C. gattii* population morphology, accompanied by increased capsule diameter and cell wall remodelling, including alterations in chitin. These changes were associated with enhanced fungal persistence within the lung. Together, these results demonstrate that *C. gattii* senses and responds to co-infecting bacterial cues, highlighting a previously underappreciated role for cross-kingdom interactions in shaping *C. gattii* adaptation and persistence within the pulmonary niche.

## Results

### C. gattii responds to mycobacterial presence by reprogramming population dynamics toward titan enrichment

Population heterogeneity, comprising titan cells (≥10 μm), regular yeast cells (5-9 μm), and small cells (≤5 μm), is a defining feature of *Cryptococcus* pathobiology^17,46^. However, the regulation and functional significance of this heterogeneity remain less well characterised for *C. gattii* in polymicrobial settings. To ensure biosafety compliance while maintaining physiological relevance, we employed heat-killed *M. tuberculosis* (HK-Mtb), which preserves mycobacterial physical components while eliminating the need for BSL-3 containment. In our co-culture experiments, *C. gattii* cells were incubated either alone or with increasing concentrations of HK-Mtb under physiological conditions (37°C, 5% CO₂), with samples collected over time for comprehensive phenotypic analysis. After 72 h of co-incubation with mycobacteria, *C. gattii*, exhibited a marked dose-dependent shift in population architecture and increase in cell body diameter (Fig. 1A-F). Consistent with our previous observations with *C. neoformans*^45^ , the proportion of titan cells increased from 6.5% in monoculture to 16.6% at the highest mycobacterial exposure accompanied by a reciprocal reduction in regular yeast cells (from 88.4 to 80.3%) and smaller cells (from 5 to 3%). Remarkably, this titan enrichment occurred despite increased total fungal cell density (Fig. 1G), diverging from the classical density-dependent constraint in which titan cell formation typically increases at lower fungal densities^47^. This finding suggests that mycobacterial cues can override normal population density controls, allowing *C. gattii* to maintain titan-enriched population structure even under high-density conditions.

**Figure 1.**
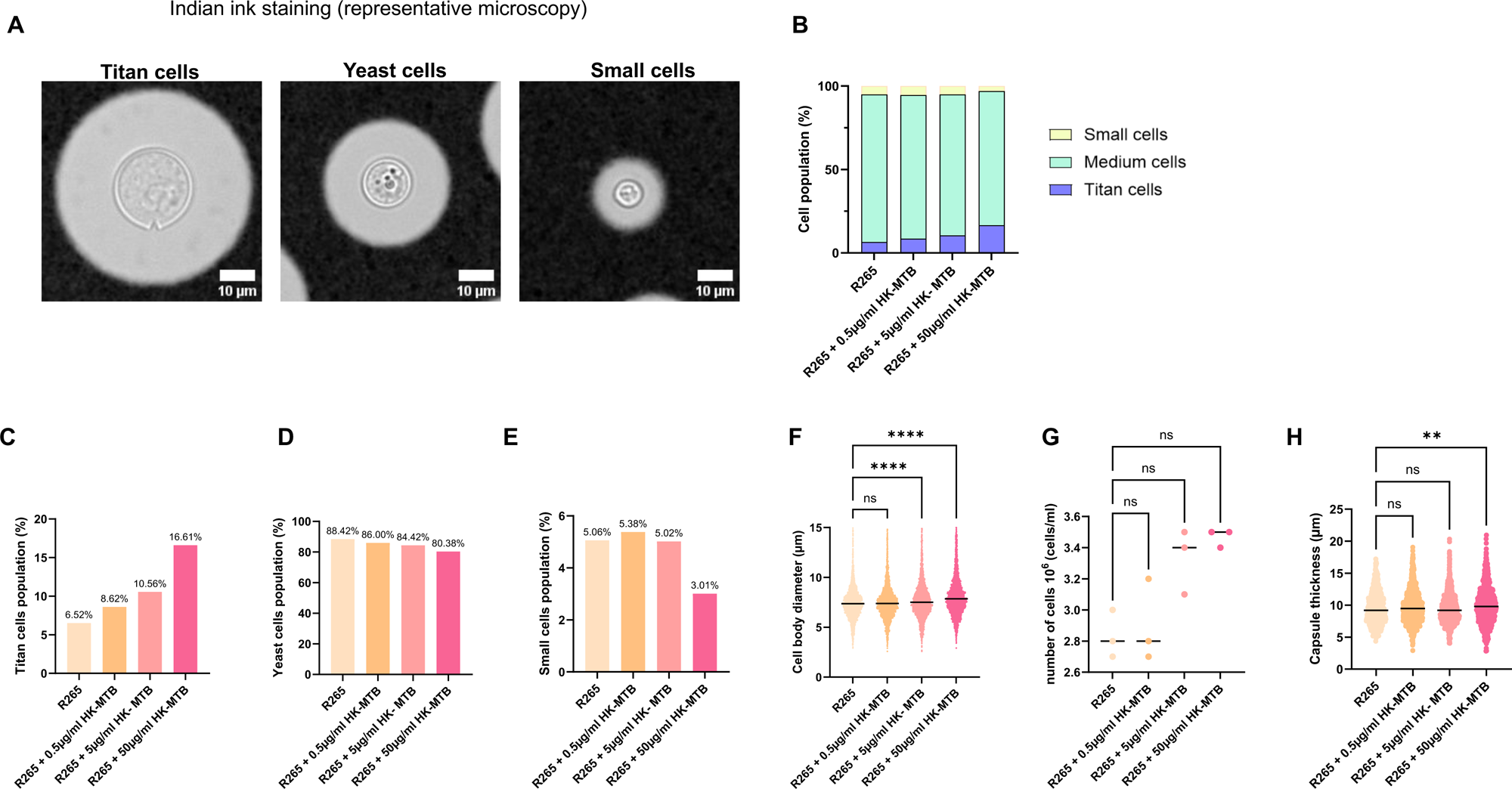
*C. gattii* responds to mycobacterial exposure by remodeling population structure toward titan enrichment. Overnight cultures of *Cryptococcus gattii* strain R265 were incubated in HPLM at 37 °C and 5% CO₂ either alone (1 × 10⁴ cells/well) or in the presence of heat-killed *Mycobacterium tuberculosis* (HK-Mtb; 0.5, 5, or 50 µg/ml) for 72 h. Cells were stained with India ink to visualise the capsule and imaged using an Olympus IX83 microscope. (A) Representative India ink microscopy images illustrating the three major morphotypes observed across conditions: titan cells (≥10 µm), yeast cells (5-9 µm), and small cells (≤5 µm). (B) Stacked bar plot summarising the proportion of each morphotype across conditions. (C-E) Quantification of the percentage of titan cells (C), yeast cells (D), and small cells (E). (F) Distribution of cell body diameter across conditions following co-incubation with increasing concentrations of HK-Mtb. (G) Total fungal cell density following exposure to HK-Mtb, measured using a Vi-CELL BLU viability analyser. (H) Capsule thickness measurements across conditions. Cell body diameter and capsule thickness were quantified using Fiji. For morphometric analyses, n = 200 cells per condition were analysed across 15 technical replicates from five biological replicates. Statistical significance was assessed using one-way ANOVA with Tukey’s multiple-comparison test. P ≤ 0.05 (*), P ≤ 0.01 (), P ≤ 0.001 (*), and P ≤ 0.0001 (****). NS indicates not significant.

### C. gattii expands capsule thickness across morphotypes in response to mycobacterial presence

The polysaccharide capsule is a central virulence determinant in *Cryptococcus*^13,14,16^, yet capsule regulation in *C. gattii* remains incompletely defined, particularly in polymicrobial environments. We analysed whether capsule architecture is modulated as part of its response to mycobacterial co-existence. When cultured in HPLM, *C. gattii* exhibited a significant increase in capsule thickness after 72 h in the presence of higher mycobacterial exposure levels (50 µg/ml) compared with monoculture controls (Fig. 1H). These data demonstrate that *C. gattii* responds to mycobacterial presence by globally enhancing capsule thickness, reinforcing a core trait linked to immune shielding and persistence^13^.

### C. gattii remodels chitin during co-culture with mycobacteria

Although the polysaccharide capsule forms the outermost fungal structure, the underlying cell wall plays a critical structural and regulatory role, influencing capsule anchoring, capsule stability, and fungal fitness^48,49^. Genetic perturbations of cell wall components have been linked to differential host immune responses and disease outcomes^48–51^. To determine whether *C. gattii* adjusts core cell wall architecture in response to mycobacterial presence, we assessed the abundance of key structural polysaccharides, chitin, which forms a central structural scaffold within the fungal cell wall^51^. In response to increasing mycobacterial exposure, *C. gattii* exhibited a consistent increase in total chitin content across the population (Fig. 2A-E). This increase was observed across all morphotypes, with titan cells (Fig 2. C), regular yeast cells (Fig 2. D), and small cells (Fig 2. E) displaying a comparable upward shift in chitin signal with increasing mycobacterial burden. The distribution profiles indicate that this response occurs broadly across the population rather than being restricted to a specific subpopulation. Together, these findings demonstrate that *C. gattii* mounts a coordinated, dose-dependent increase in chitin abundance in response to mycobacterial presence.

**Figure 2.**
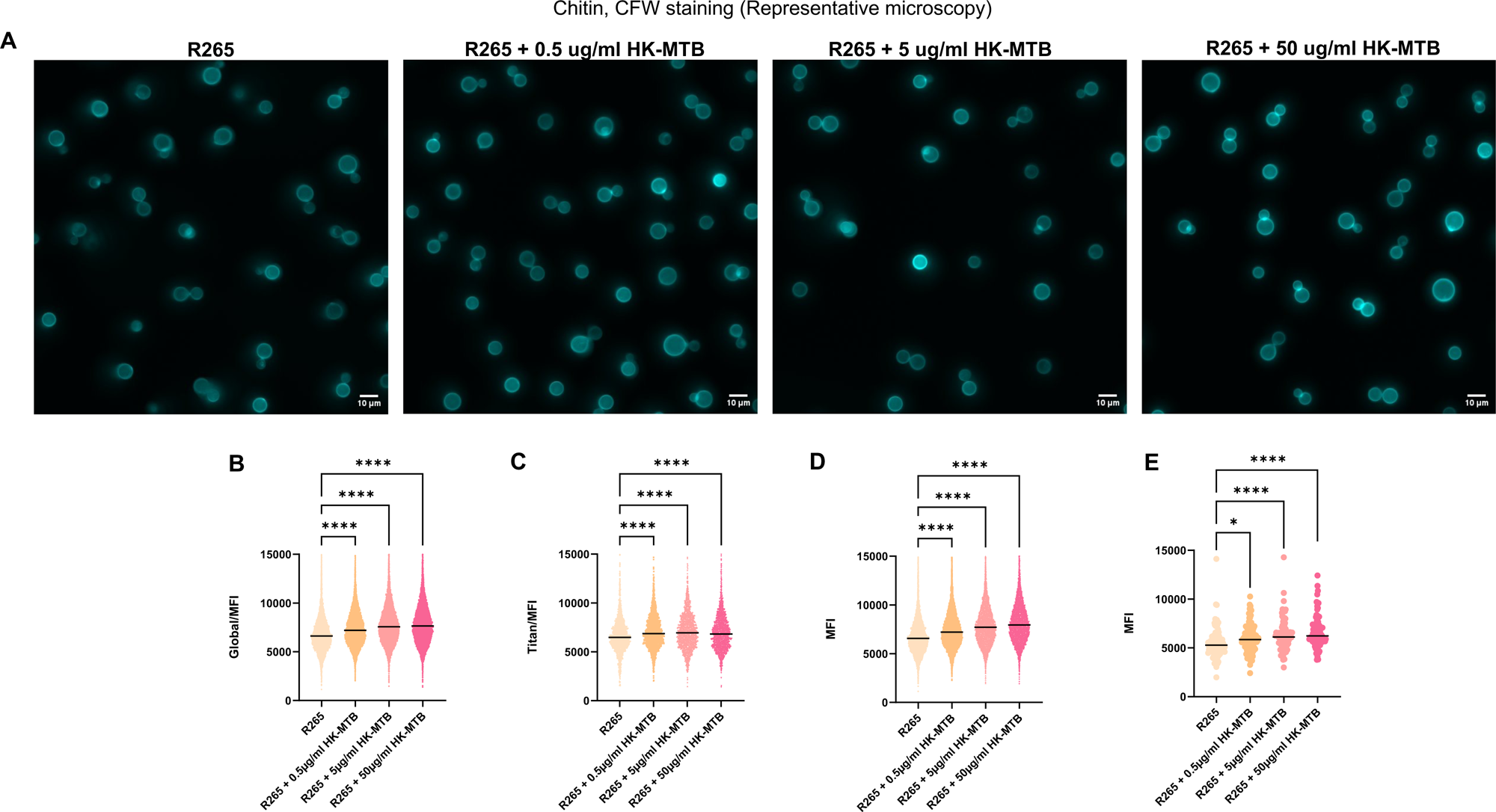
*C. gattii* remodels chitin during co-culture with mycobacteria. *Cryptococcus gattii* strain R265 was incubated in human plasma-like medium (HPLM) at 37 °C and 5% CO₂ either alone or in the presence of increasing concentrations of heat-killed *Mycobacterium tuberculosis* (HK-Mtb; 0.5, 5, and 50 µg/ml) for 72 h. Chitin was visualised using Calcofluor White (CFW) staining and quantified by fluorescence intensity. (A) Representative fluorescence microscopy images of CFW-stained cells across conditions. (B) Global chitin fluorescence intensity measured across the total fungal population. (C-E) Quantification of chitin abundance within distinct morphological subpopulations, including (C) titan cells (≥10 µm), (D) yeast cells (5-9 µm), and (E) small cells (≤5 µm). Fluorescence intensity was quantified using FIJI software and analysed in GraphPad Prism. Data are presented as single-cell fluorescence intensity distributions from five biological replicates. Morphological categories were defined according to cell body diameter (titan ≥10 µm; yeast 5-9 µm; small ≤5 µm). Statistical significance between conditions was assessed using Kruskal-Wallis analysis with multiple comparisons. P ≤ 0.05 (*), P ≤ 0.0001 (*\*\*\*), and ns indicates not significant.

### Chitosan remodelling in C. gattii shows biphasic response to mycobacteria

A closely linked chitin metabolism byproduct, chitosan, contributes to cell wall flexibility, stress resistance, and capsule attachment^51^. At lower levels of mycobacterial exposure, *C. gattii* increased chitosan abundance across yeast cells, and regular yeast cells (Fig. 3A-E). In contrast, higher mycobacterial exposure was associated with a marked reduction in chitosan levels across titan cells (Fig 3. C), regular yeast cells (Fig 3. D), and small cells (Fig 3. E). This biphasic response indicates that *C. gattii* independently regulates chitosan levels rather than passively reflecting upstream changes in chitin synthesis, suggesting a finely tuned, dose-dependent adjustment of chitosan in response to mycobacterial cues. Importantly, this coordinated increase in chitin and biphasic modulation of chitosan was recapitulated in the independent clinical isolates WM779 and M27055. These isolates were isolated from animals and human which were isolated from veterinary and human respectively^52,53^ (Supplementary Fig. 1 and 2), supporting that this adaptive cell wall remodelling response is conserved across *C. gattii* strains

**Figure 3.**
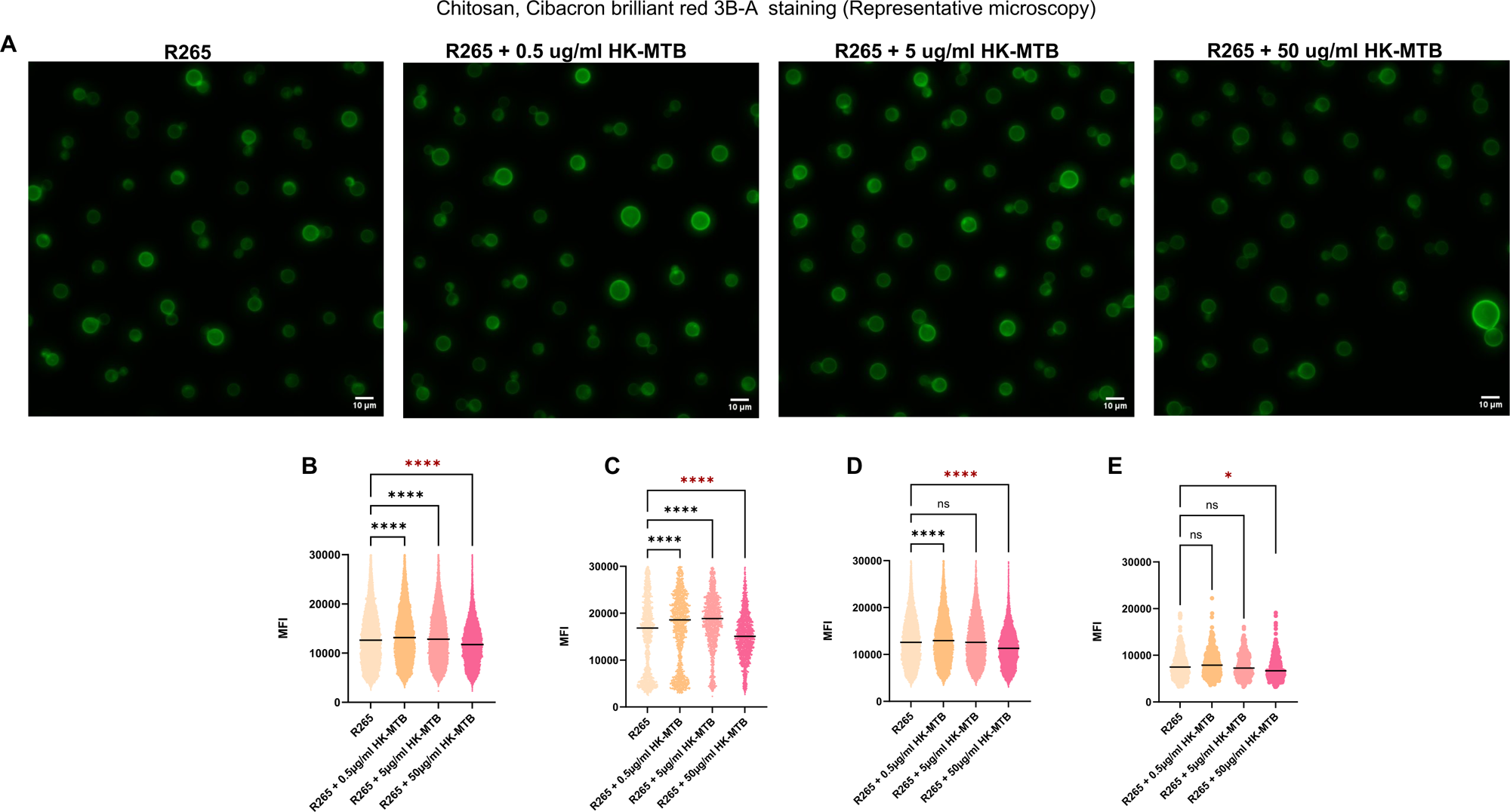
Chitosan remodelling in *C. gattii* shows biphasic response to mycobacteria. *Cryptococcus gattii* strain R265 was incubated in human plasma-like medium (HPLM) at 37 °C and 5% CO₂ either alone or in the presence of increasing concentrations of heat-killed *Mycobacterium tuberculosis* (HK-Mtb; 0.5, 5, and 50 µg/ml) for 72 h. Chitosan was visualised using Cibacron Brilliant Red 3B-A staining and quantified as mean fluorescence intensity (MFI). (A) Representative fluorescence microscopy images of Cibacron Red-stained cells across conditions. (B) Global chitosan fluorescence intensity measured across the total fungal population. (C-E) Quantification of chitosan abundance within distinct morphological subpopulations, including (C) titan cells (≥10 µm), (D) yeast cells (5-9 µm), and (E) small cells (≤5 µm). Fluorescence intensity was quantified using FIJI software and analysed in GraphPad Prism. Data are presented as single-cell fluorescence intensity distributions from five biological replicates. Morphological categories were defined according to cell body diameter (titan ≥10 µm; yeast 5-9 µm; small ≤5 µm). Statistical significance between conditions was assessed using Kruskal-Wallis analysis with multiple comparisons. P ≤ 0.05 (\**), P ≤ 0.0001 (*\*\*\*), and ns indicates not significant.

### C. gattii exhibits enhanced pulmonary persistence in vivo following mycobacterial exposure

*C. gattii* establishes prolonged infection within the lung, yet the factors that promote pulmonary persistence *in vivo* remain incompletely defined. Here we wanted to test if the observed changes *in vitro* increased Titan population, increased capsule, and cell wall remodelling during co-culture with mycobacteria have impact on the outcome of infection *in vivo*. We used a murine inhalation model of *C. gattii* infection to establish infection, followed by intranasal inoculation with HK-Mtb 24 h later. Mice exposed to HK-Mtb exhibited increased *C. gattii* colony-forming units (CFUs) in the lung compared with animals infected with *C. gattii* alone, 72 h post infection (Fig. 4). This data suggests that *M. tuberculosis* co-infection may promote *C. gattii* growth or survival within the pulmonary environment, providing *in vivo* evidence that fungal responses observed *in vitro* are associated with enhanced pulmonary fitness.

**Figure 4.**
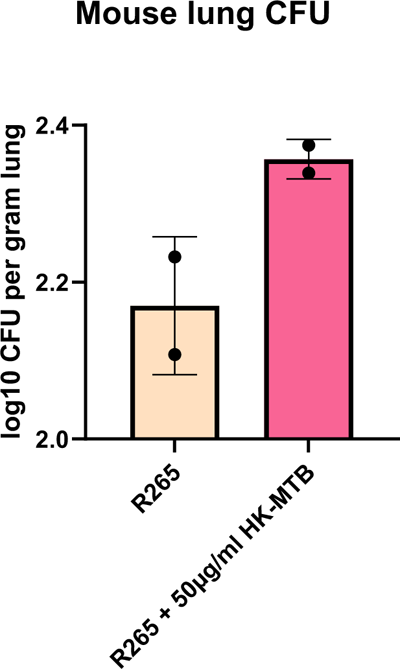
Mycobacterial exposure enhances pulmonary persistence of *Cryptococcus gattii in vivo*. Two groups of female C57BL/6 8-12 weeks mice were infected intranasally with *Cryptococcus gattii* strain R265 (50 yeast cells/mouse), and 24 hour later one exposed to 50 µg ml⁻¹ HK-Mtb in 50 ul Phosphate free buffer. At 3 days post infection, mice were euthanized and lungs were homogenized in sterile PBS, and serial dilutions were plated on YPD agar supplemented with 1% penicillin-streptomycin. Plates were incubated at 30 °C for 48 h, after which colony-forming units (CFU) were enumerated to determine fungal burden. Fungal burden is presented as log₁₀ CFU per lung. N=2 mice per group.

Overall, our findings reveal that across population structure, capsule architecture, cell wall composition, and *in vivo* persistence, *C. gattii* responds to mycobacterial presence through an integrated adaptive programme. These responses favour persistence-associated morphotypes, reinforce capsule organisation, and remodel structural polymers, collectively supporting fungal fitness in response to mycobacteria, with potential implications for pathogenesis and disease outcome in patients carrying a co-infection.

## Discussion

Fungal pathogens rarely exist in isolation within their natural or host-associated environments, yet the consequences of polymicrobial coexistence for fungal biology and disease outcome remain poorly understood. This gap is clinically significant: in high-burden settings, co-infection with multiple pulmonary pathogens is the rule rather than the exception, yet treatment strategies are almost universally designed around single-pathogen paradigms. Previous studies have demonstrated that bacterial interactions can directly modulate canonical cryptococcal virulence traits^30,31,35,45^. For example, *Klebsiella aerogenes* enhances melanisation of *C. neoformans* through the production of melanin precursors, whereas other bacterial signals suppress melanin synthesis, highlighting how microbial cues influence pigment production, a factor linked to stress resistance and virulence^54^. Consistent with these observations, bacterial perturbation of the microbial environment can alter titan cell formation in *Cryptococcus*^30,31^. The present study extends this framework to a clinically critical pairing, *C. gattii* and *Mycobacterium tuberculosis*, two co-endemic pulmonary pathogens whose interaction at the biological level has not previously been characterised.

In this study, we demonstrate that *C. gattii* responds to mycobacterial-derived cues through a coordinated programme of population restructuring, capsule expansion, cell wall remodelling, and enhanced pulmonary persistence *in vivo*. Together, these findings indicate that *C. gattii* does not simply tolerate the presence of *M. tuberculosis*-derived signals but actively interprets and responds to them in ways that favour fungal fitness within a shared pulmonary niche. A central finding of this work is that *C. gattii* reprograms population dynamics toward titan cell enrichment during mycobacterial co-culture. Population heterogeneity is increasingly recognised as a key virulence strategy in *Cryptococcus*, yet most mechanistic insight has been derived from *C. neoformans*^17,46^. Our data show that *C. gattii*, a species with distinct epidemiology, host range, and disease presentation^55,56^ , similarly exhibits morphological plasticity and regulated population restructuring. Notably, titan enrichment occurred despite increasing fungal density, a condition under which titan cell frequency typically declines in conventional culture, suggesting that titan formation can be driven independently of classical density-associated quorum constraints when external microbial signals are present^17,47^. These findings indicate that population architecture in *C. gattii* is not governed solely by intrinsic quorum-sensing mechanisms but may also be shaped by environmental signals encountered within polymicrobial settings, a regulatory flexibility that warrants further mechanistic investigation.

Capsule expansion represents another major adaptive response. The polysaccharide capsule is one of the principal virulence determinants in *Cryptococcus*^13,14,16^, yet its regulation in *C. gattii* remains comparatively understudied relative to *C. neoformans*, and the environmental signals governing it are incompletely defined. Our observation that capsule thickness increased across the population rather than being confined to a discrete enlarged subpopulation suggests that *C. gattii* engages capsule expansion as a coordinated adaptive response to the polymicrobial environment, rather than this arising solely from selective enrichment of pre-existing heavily encapsulated cells. This interpretation is supported by the broad distributional shift in capsule measurements across the population, consistent with an active stress- or pathogen-sensing response. Given the central role of the capsule in complement evasion, resistance to oxidative and nitrosative killing, and intracellular survival following macrophage phagocytosis^13,14,16^, this response is likely to confer substantial survival advantage within the inflamed pulmonary environment that characterises TB disease.

Beyond capsule architecture, our data reveal structured, polymer-specific remodelling of the cell wall during mycobacterial exposure. Importantly, these changes were not uniform: chitin and chitosan displayed dose-dependent and morphotype-specific modulation. Although the cell wall lies beneath the capsule, its composition strongly influences capsule anchoring, capsule dimensions, and the surface mechanical properties that govern immune recognition. Previous genetic studies have demonstrated that perturbations in chitin and chitosan biosynthesis alter capsule organisation, stress tolerance, and virulence in *Cryptococcus*^48,49,51^. Notably, chitosan, through its cationic charge, is thought to mask immunostimulatory cell wall components, thereby dampening host pattern recognition; its modulation during mycobacterial exposure suggests that *C. gattii* may additionally be tuning its immune visibility in polymicrobial contexts. Taken together, the polymer-specific remodelling observed here suggests that *C. gattii* fine-tunes internal structural layers to stabilise capsule expansion and maintain surface integrity when a co-resident pulmonary pathogen is present.

Exposure to heat-killed mycobacteria in a murine inhalation model was associated with increased pulmonary burden of *C. gattii*, consistent with increased number of cells *in vitro*, supporting the hypothesis that mycobacterial-derived signals can directly enhance fungal persistence within the lung, independently of active mycobacterial replication or the immune dysregulation it causes. Although non-replicative, heat-killed mycobacteria retain structurally intact cell wall components, including mycolic acids, lipoarabinomannan, and peptidoglycan, capable of engaging both host and microbial pattern recognition and sensing pathways. The fact that these acellular cues are sufficient to elicit robust fungal phenotypic adaptation and enhanced *in vivo* survival strongly suggests that *C. gattii* harbours dedicated sensing mechanisms capable of detecting and interpreting signals from co-resident bacterial pathogens, rather than benefiting indirectly from immune suppression driven by live infection.

Our results align with a growing body of evidence indicating that fungi actively respond to neighbouring microbes rather than passively experiencing their effects. In *Candida albicans*, direct interaction with bacterial species can trigger morphological switching, biofilm formation, and altered cell wall architecture^37,57^. In *Aspergillus fumigatus*, bacterial metabolites influence hyphal growth, secondary metabolism, and stress responses^58,59^. The present findings extend this framework to a co-infection dyad of major global health importance: *C. gattii* and *M. tuberculosis* together account for substantial morbidity in overlapping geographic regions, yet their biological interaction has remained entirely unexplored. Together, these findings support the emerging concept that polymicrobial interactions are not merely ecological by-products of co-infection but biologically meaningful processes capable of reshaping pathogen behaviour, fitness, and disease progression within the host, and that this reshaping is likely to have direct consequences for treatment response and clinical outcome.

The implications of these findings are particularly significant for *C. gattii*, and may help illuminate one of the most perplexing unresolved questions in medical mycology: why this organism causes severe disease in people with apparently intact immunity. Unlike *C. neoformans*, which overwhelmingly affects severely immunocompromised individuals, particularly those with advanced HIV disease, *C. gattii* has a substantially broader host range. A striking proportion of *C. gattii* infections occur in individuals without detectable immunosuppression, yet the pathogen still causes severe, treatment-refractory disease across the immune spectrum. This capacity to breach intact immune defences distinguishes *C. gattii* from most other medically important fungi and remains mechanistically unexplained; proposed explanations, including subtle undetected immune defects, heightened intrinsic virulence, or structural differences in capsule composition, remain unproven.

*C. gattii* is also notable as a true outbreak pathogen, a designation rare among fungi. The landmark Vancouver Island outbreak beginning in 1999, driven by the hypervirulent VGII molecular type, demonstrated that *C. gattii* can emerge *de novo* in previously non-endemic temperate regions and sustain environmental transmission to immunocompetent hosts over years^60^. This outbreak, and subsequent spread throughout the Pacific Northwest, revealed a capacity for geographic range expansion now increasingly linked to climate-driven shifts in fungal ecology, raising the prospect of future emergence in regions currently considered non-endemic^60^. The outbreak strain’s enhanced virulence has been attributed in part to differences in capsule structure, titan cell induction, and macrophage evasion, yet the precise adaptations conferring epidemic potential remain unresolved. *C. gattii* is further characterised by prominent pulmonary involvement, a propensity for cryptococcoma formation, and comparatively lower rates of CNS dissemination than *C. neoformans*. How *C. gattii* persists within the lung for prolonged periods, sometimes years before clinical presentation, and what determines the transition to extrapulmonary dissemination, are critical unanswered questions. Our data suggest that microbial coexistence within the lung may represent an underappreciated driver of this persistence, dynamically reshaping population structure and surface architecture in ways that favour immune evasion and survival. For *C. gattii* specifically, where the lung is both the primary infection site and the battleground on which disease outcome is determined, understanding how co-resident bacterial pathogens shape fungal adaptation may be essential to explaining its unusual virulence in immunocompetent hosts and the conditions that enabled an epidemic strain to emerge.

Several important questions remain: the molecular mechanisms by which *C. gattii* senses mycobacterial-derived cues are currently unknown. Future research should investigate potential sensing pathways may involve direct recognition of mycobacterial cell wall components, such as mycolic acids or lipoarabinomannan, by fungal pattern recognition or stress-sensing systems, or indirect sensing of host-derived inflammatory mediators induced by mycobacterial products. Dissecting these pathways, for instance through transcriptomic and proteomic profiling of *C. gattii* responses to defined mycobacterial fractions, will be essential to understanding how fungal decision-making is integrated within polymicrobial environments. In addition, while heat-killed mycobacteria were sufficient to elicit robust fungal responses *in vitro* and enhanced pulmonary persistence *in vivo*, future studies using live co-infection models will be required to capture the full complexity of fungal-mycobacterial-host tripartite interactions, including the contribution of active immune modulation by viable MTb. Ultimately, understanding these sensing and adaptive mechanisms has direct therapeutic relevance. If mycobacterial signals drive *C. gattii* toward enhanced persistence and immune evasion, then disrupting this cross-kingdom signalling or the downstream adaptive programmes it activates could represent a tractable target for adjunctive host-directed or anti-virulence therapy. Such strategies would be particularly valuable in co-endemic settings where TB and cryptococcal disease overlap, and where current treatment algorithms do not account for the biological consequences of pathogen coexistence.

## Materials and Methods

### Strains and culture conditions

*Cryptococcus gattii* and *Mycobacterium tuberculosis* strains used in this study are summarized in [Fig. S1/Table X, Supplementary Material]. The *C. gattii* strain R265, 779, 27055 (VGII) and the heat killed *M. tuberculosis* clinical isolate were kindly provided by [Farrar and Brown laboratories, University of Exeter]. Yeast cells were routinely maintained on yeast extract peptone dextrose (YPD) agar plates and stored at 4 °C. For routine culture, cells were inoculated into 5 ml YPD broth in 50 ml Falcon tubes and incubated overnight (12-16 h) at 30 °C with shaking at 200 r.p.m. Following overnight growth, cell density was determined using a Vi-CELL BLU viability analyser (Beckman Coulter) and adjusted in phosphate-buffered saline (PBS; pH 7.4) for subsequent experiments.

HK-Mtb was suspended in HPLM supplemented with penicillin-streptomycin to maintain sterility during co-incubation assays.

For co-culture experiments, *C. gattii* cells were incubated either alone or in the presence of increasing concentrations of HK-Mtb. Cultures were maintained at 37 °C with 5% CO₂, and samples were collected over the course of the experiment for downstream analyses. Each condition was performed in triplicate, and experiments were repeated across five independent biological replicates.

### Quantification of Cryptococcus gattii Cell Density

Total *Cryptococcus gattii* cell density was quantified following 72 h co-incubation with HK-Mtb. Samples were diluted 1:100 in PBS and analysed using a Vi-CELL BLU automated cell viability analyser (Beckman Coulter) to determine total cell concentration (cells/ml). Measurements were performed for *C. gattii* cultured alone or co-incubated with 0.5, 5, or 50 µg/ml HK-Mtb. Data from six independent biological replicates were compiled and analysed using GraphPad Prism to assess the effect of mycobacterial exposure on total fungal cell density.

### India ink staining and image acquisition

India ink staining was used to visualise the *Cryptococcus gattii* cell body and capsule. Briefly, 3 μl of cell suspension was mixed with 3 μL India ink on a glass slide and imaged using an Olympus IX83 microscope under identical acquisition settings across conditions. Samples included *C. gattii* cultured alone or co-incubated with 0.5, 5, or 50 µg/ml HK-Mtb. Images were acquired and processed using ImageJ2 (Fiji).

### Measurement of C. gattii cell body diameter and capsule thickness

India ink stain was used to quantify cell body diameter and capsule thickness. Images were analysed in Fiji, and cells were randomly selected for measurement. At least 400 cells per condition were quantified. Measurements were exported for analysis in GraphPad Prism. Cells were categorised according to cell body diameter as titan cells (≥10 μm), yeast cells (≥5 μm and <10 μm), and small cells (<5 μm) as previously described.

### Fluorescence Microscopy Analysis of Cryptococcus gattii Cell Wall Components

Cell wall composition of *Cryptococcus gattii* strain R265 was analysed following exposure to 0.5, 5, or 50 µg/ml HK-Mtb. The abundance of chitin, and chitosan, was assessed using fluorescent staining. Cells were adjusted to 5.0 × 10^6^ cells/ml of suspension was used for each staining condition. After staining, cells were collected by centrifugation, washed twice with PBS, and resuspended in 500 μl PBS. A 20 μl aliquot of the stained suspension was transferred to ibidi 8-well μ-slides for imaging. Samples included *C. gattii* cultured alone or co-incubated with 0.5, 5, or 50 µg/ml HK-Mtb. Fluorescence images were acquired using an Olympus IX83 fluorescence microscope with constant exposure settings across all conditions. Fluorescence intensity was quantified using FIJI (ImageJ) and analysed using GraphPad Prism.

### Chitin staining

Chitin was visualised using Calcofluor White (CFW) at a final concentration of 5 μg/ml in PBS (pH 7.2). Cells were incubated for 15 min at room temperature in the dark, washed with PBS, and imaged using the DAPI channel.

### Chitosan staining

Chitosan was detected using Cibacron Red at a final concentration of 10 μg/ml in PBS adjusted to pH 2. Cells were incubated for 20 min at room temperature in the dark, washed with PBS, and imaged using the mCherry channel.

### Statistical analysis

Data were analysed using GraphPad Prism 10.6.1 (892) with ANOVA. Medium with interquartile range is analysed using the Kruskal-Wallis Test, and Spearman’s coefficient for p-value. The target p-value thresholds include p<0.05, p<0.01, p<0.001, and p<0.0001.

## Acknowledgement

We thank the staff of the animal facilities at the University of Exeter for the care and support of our animals. We acknowledge funding from the MRC Centre for Medical Mycology at the University of Exeter (MR/N006364/2 and MR/V033417/1), the NIHR Exeter Biomedical Research Centre (NIHR203320), and the MRC Doctoral Training Grant MR/W502649/1. H.I. is supported by the MRC Doctoral training Grant MR/W502649/1. The views expressed are those of the author(s) and not necessarily those of the NIHR or the Department of Health and Social Care. For the purpose of open access, the author has applied a CC BY public copyright licence to any Author Accepted Manuscript version arising from this submission.

---

**Supplementary Figure 1.**
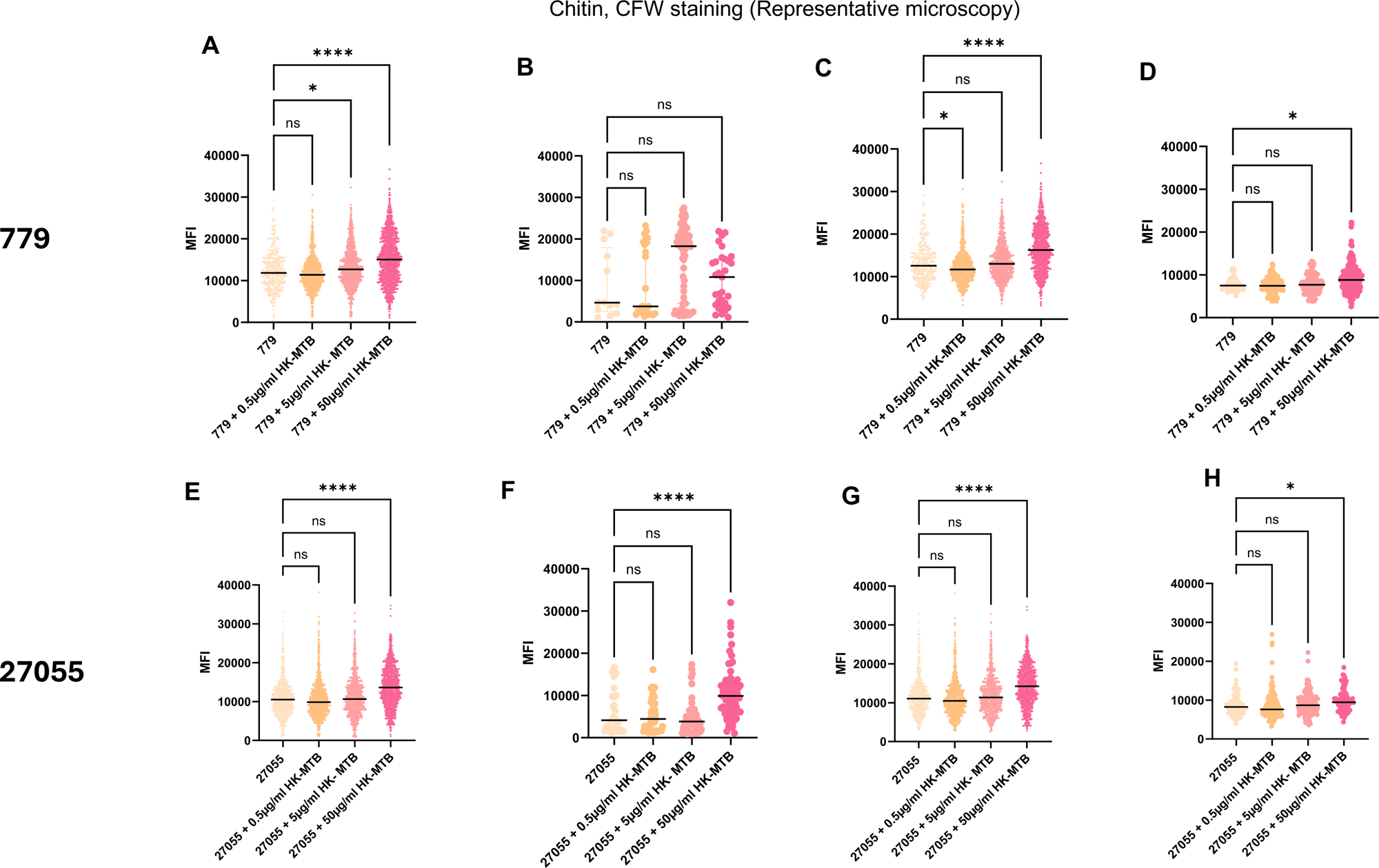
Conserved chitin remodelling across *C. gattii* clinical isolates following mycobacterial exposure. Additional *Cryptococcus gattii* isolates WM779 and M27055 were incubated in human plasma-like medium (HPLM) at 37 °C and 5% CO₂ either alone or in the presence of increasing concentrations of heat-killed *Mycobacterium tuberculosis* (HK-Mtb; 0.5, 5, and 50 µg/ml) for 72 h. Chitin was visualised using Calcofluor White (CFW) staining and quantified as mean fluorescence intensity (MFI). (A-D) Quantification of chitin fluorescence intensity across WM779 morphological subpopulations, including (A) global population, (B) titan cells (≥10 µm), (C) yeast cells (5-9 µm), and (D) small cells (≤5 µm). (E-H) Quantification of chitin fluorescence intensity across M27055 morphological subpopulations, including (E) global population, (F) titan cells (≥10 µm), (G) yeast cells (5-9 µm), and (H) small cells (≤5 µm). Fluorescence intensity was quantified using FIJI software and analysed in GraphPad Prism. Data are presented as single-cell fluorescence intensity distributions from five biological replicates. Morphological categories were defined according to cell body diameter (titan ≥10 µm; yeast 5-9 µm; small ≤5 µm). Statistical significance between conditions was assessed using Kruskal-Wallis analysis with multiple comparisons. P ≤ 0.05 (\**), P ≤ 0.0001 (*\*\*\*), and ns indicates not significant.

**Supplementary Figure 2.**
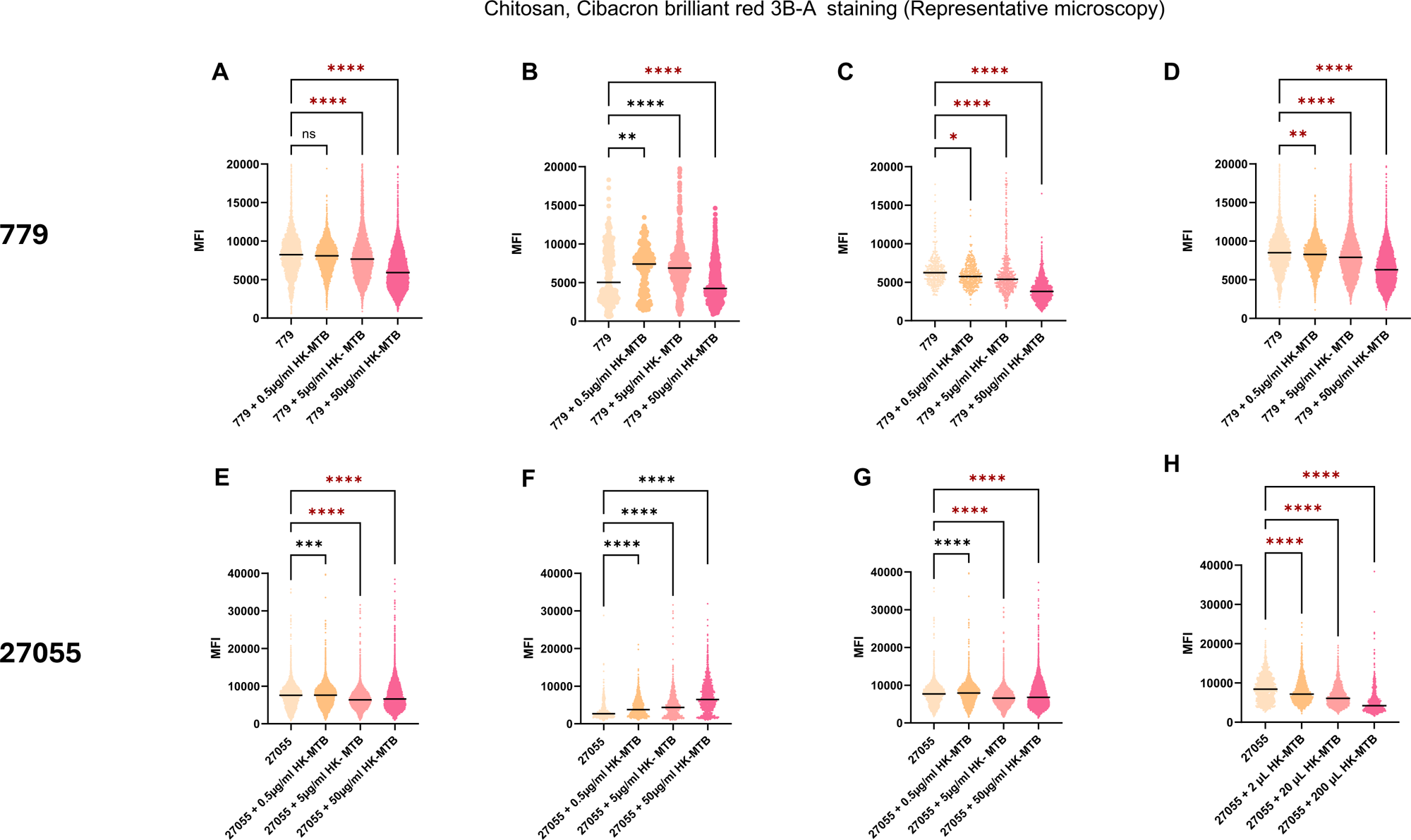
Conserved biphasic chitosan remodelling across distinct *C. gattii* isolates following mycobacterial exposure. Additional *Cryptococcus gattii* isolates WM779 and M27055 were incubated in human plasma-like medium (HPLM) at 37 °C and 5% CO₂ either alone or in the presence of increasing concentrations of heat-killed *Mycobacterium tuberculosis* (HK-Mtb; 0.5, 5, and 50 µg/ml) for 72 h. Chitosan was visualised using Cibacron Brilliant Red 3B-A staining and quantified as mean fluorescence intensity (MFI). (A-D) Quantification of chitosan fluorescence intensity across WM779 morphological subpopulations, including (A) global population, (B) titan cells (≥10 µm), (C) yeast cells (5-9 µm), and (D) small cells (≤5 µm). (E-H) Quantification of chitosan fluorescence intensity across M27055 morphological subpopulations, including (E) global population, (F) titan cells (≥10 µm), (G) yeast cells (5-9 µm), and (H) small cells (≤5 µm). Fluorescence intensity was quantified using FIJI software and analysed in GraphPad Prism. Data are presented as single-cell fluorescence intensity distributions from five biological replicates. Morphological categories were defined according to cell body diameter (titan ≥10 µm; yeast 5-9 µm; small ≤5 µm). Statistical significance between conditions was assessed using Kruskal-Wallis analysis with multiple comparisons. P ≤ 0.05 (\**), P ≤ 0.0001 (*\*\*\*), and ns indicates not significant.

